# Green synthesis, characterization, and potential antimicrobial studies of Ag-MgO nanocomposites mediated from *Talinum triangulare* leaf extract

**DOI:** 10.1101/2024.11.08.622651

**Authors:** Fonye Nyuyfoni Gildas, Paboudam Gbambie Awawou, Kwati Leonard, Gilbert Njowir Ndzeidze, Francois Eya’ane Meva

**Author notes:** Correspondingauthors: Paboudam Gbambie Awawou,; Francois Eya’ane Meva.

## Abstract

In this study, Ag-MgONCs (silver-magnesium oxide nanocomposites) were fabricated using *Talinum triangulare* leaf extract as a renewable, mild reducing, and stabilizing agent. The synthesis process involved molecular interactions between pre-prepared AgNPs (silver nanoparticles) and *in situ*-prepared MgO. Confirmation of the nanocomposites came from micrographic images obtained through SEM-EDX analysis. Dynamic light scattering (DLS) revealed a characteristic nanocomposite size of 295 nm. The infrared spectrum (FTIR) proved interactions between *T. triangulare* and Ag. Additionally, X-ray patterns of AgNPs and Ag-MgONCs confirmed hexagonal and cubic crystalline states, respectively, with average particle sizes of 17 nm. Furthermore, an *in vitro* analysis against five bacterial strains (*Escherichia coli*, *Acinetobacter baumannii*, *Pseudomonas aeruginosa*, *Klebsiella pneumoniae*, and *Proteus mirabilis*) revealed pronounced activity of the nanocomposites, particularly against *Proteus mirabilis*, indicating bactericidal potential.

## 1. INTRODUCTION

Nanoparticles, typically ranging from 1 to 100 nm, exhibit vastly superior properties compared to their larger bulk material counterparts [1]. A composite is a material composed of several components, and which is called nanocomposite at the nanometric scale. They have broad consideration because of their prospects to combine fascinating properties of different nanoscale systems to upsurge optical, electronic, magnetic, and mechanical properties. Nanocomposites possess a matrix, in which different materials are combined to develop a material with new properties i.e., one or more nanosized materials embedded either in a ceramic, metal, or polymer matrix connected by weak electrostatic interactions (Van der Waals, hydrogen bonding) or by covalent bonds. It consists of two parts, a continuous phase, and a discontinuous reinforcing phase. Thus, a nanocomposite is a multiphase with at least one phase on the nanoscale dimension whichhas a combination or has markedly different properties from the component materials.

Silver nanoparticles are undoubtedly the most widely used inorganic nanoparticles with tremendous applications in the field of sensitive biomolecular detection and antibacterial activities [2]. MgO, on the other hand, has recently received greater attention due to its vast applications in biomedical materials, catalysis, and as absorbents [3]. However, based on the antibacterial properties of MgO having a less potent bactericidal effect compared to silver-based antibacterial agents, efforts are now being channeled into the development of composite materials to exert strong synergistic antibacterial activity against infectious pathogens through chemical methods [4]. However, the use of nanosynthesized biomedical materials has been a major concern due to their high cost, toxic nature, and production of nonecofriendly by-products and nonbiodegradable stabilizing agents posing a danger to the environment and humans [5].

In the literature, this shortcoming has been addressed by successive biosynthetic approaches using plant extracts have been explored and reported [6]. Plant-based approaches have many advantages over chemical methods. According to previous investigations, the polyol components and the water-soluble heterocyclic components in these plants play a significant role in the reduction of metallic ions. They have proven to be effective capping and stabilizing agents for nanoparticles with widespread technological and medicinal applications [7]. The use of the extract of differentplant parts has been extensively reported and reviewed for silver nanoparticles which have been successfully synthesized through different plant species extract.

*Talinum triangulare* (water leaf) is an herbaceous perennial, calescent, and glabrous plant that is widely grown in tropical regions as a leaf vegetable with high water content. African leafy vegetables are known for their nutritive content and their prophylactic and therapeutic properties due to the presence of bioactive compounds like carbohydrates, amino acids, proteins, lipids, and antioxidants. *T. triangulare* is consumed as a vegetable and sauce constituent. Nutritionally, it has been shown to possess essential nutrients such as b-carotene, minerals (such as calcium, potassium, and magnesium), pectins, proteins, and vitamins [8]. It has also been implicated in the management of cardiovascular diseases such as stroke and obesity [9]. According to phytochemical analysis carried out by Aja et al. on the leaf extract of *T. fructicosum*, an appreciable amount of secondary metabolites like flavonoids, alkaloids, saponins, and low levels of toxicants like tannins were found [10]. This plant has also been established as a suitable precursor for the green synthesis of potentially bioactive metal nanoparticles [11].

In this study, Ag-MgO nanocomposites will be synthesized and optimized. This will be done by varying pH, incubation conditions, extract, and silver ion quantities. The formation of the Ag-MgONCs will be confirmed by the observation of a band surface plasmon resonance in the wavelength interval of 350 to 500 nm, whose stability will be determined by aging. The latter will be characterized and its viability elucidated.

With the efforts to reduce generated hazardous waste, green synthesis and chemical processes are progressively integrating with modern developments in science and industry. Researchers are exploiting the introduction of ecofriendly materials to develop novel improved antibacterial nanomaterials against multidrug resistant human pathogens like *Escherichia coli*which is a deadly contaminants in drinking water.

## 2. MATERIALS AND METHODS

### 2.1. Selection of Bacteria strands

Escherichia coli, Acinetobacter baumannii, Pseudomonas aeruginosa, Klebsiella pneumoniae, and Proteus mirabilis were grown and test in the bacteriology unit, at the Department of Pharmaceutical Sciences, Faculty of Medicine, and University of Douala, Cameroon.

*Plant Extract.* Fresh Water leaf *(Talinum fructicosum)* was harvested on Sunday, March 24^th^, 2021 at Mvog-Betsi, Centre region, Cameroon and authenticated at the National Herbarium of Cameroon, in comparison with a voucher specimen previously deposited (Herbier 42977 HNC) (Figure 1).

**Figure 1:**
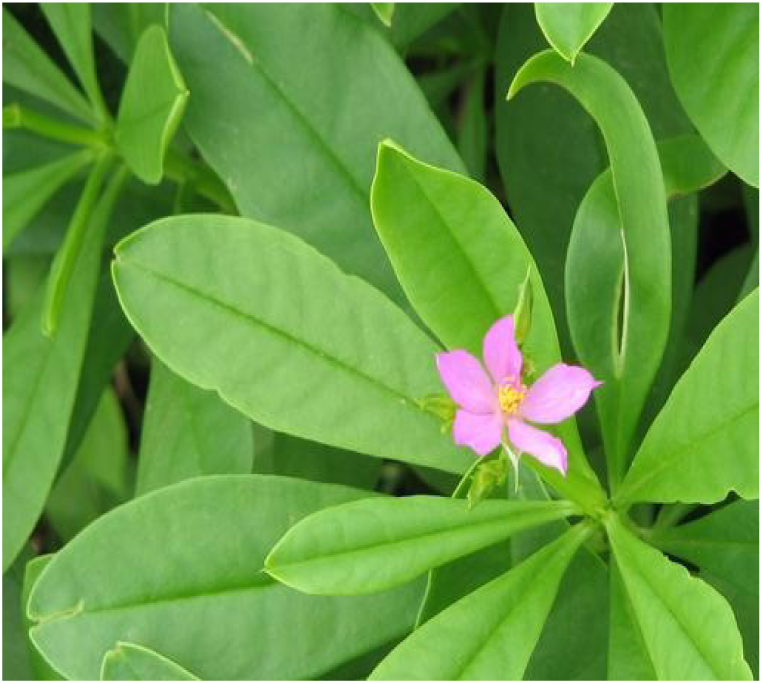
*Talinum* t*r*iangulare

*2.1. Talinum triangulare* leaves were surface cleaned with running tap water followed by deionized water to remove all the dust and unwanted visible particles. The aqueous extract of *Talinum triangulare* was prepared by boiling 25 g of finely cut leaves in 250 mL deionized water for 5 min at 80 °C and stirred for 5 min using a hot plate equipped with magnetic stirrer. The extract was filtered through Whatman No. 1 filter paper and cotton to remove particulate matter, the brown solution obtained was stored at 4 °C for further use, being usable for 1 week due to the gradual loss of plant extract viability for prolonged storage.

### 2.2. Synthesis of Silver Nanoparticles

Silver nitrate (AgNO_3_) was obtained from Sigma-Aldrich Chemicals Germany. Deionized water was used throughout the reactions. All glassware was washed with dilute nitric acid (HNO_3_) and deionized water and then dried in a hot air oven. The different experiments were done for three extract quantities named (3 mL, 4 mL and 5 mL), using 50 mL of different concentrations of AgNO_3_ (5×10^-3^ M, 5×10^-2^ M, 5×10^-1^ M) at room temperature and pressure. The resulting solutions were 1 min hand shaken and wrapped with aluminum foil paper and then incubated in the dark to minimize the photoactivation of silver nitrate. Reactions were performed under static conditions. The color of the solutions changed from an initial light brown (which is the color of the *Talinum triangulare* leaf extract) to pale brown in 5 minutes. The first hour of reaction was monitored by UV-Vis spectroscopy of 2.5 mL of the reaction suspension using an UV-visible (Ocean Insight) spectrophotometer operating at 1 nm resolution with optical length of 10 mm. Concentrations were determined by centrifugation after 24 hours of incubation. UV-Vis analysis of the reaction mixture was observed for a period of 300 s. Powder X-ray diffraction measurements were carried out using a Thermos Scientific Equinox 100 instrument by preparing a thin film of the silver-organic nanopowder on silicon substrate.

### 2.3. Synthesis of Silver Magnesium Oxide Nanocomposites

1 M Mg(OCH_2_CH_3_)_2_ was combined with 1 M Ethanol for approximately 30 minutes(Equation 1). The resulting biosynthesized Ag nanoparticles were then mixed and subsequently incorporated into the prepared MgO sol.

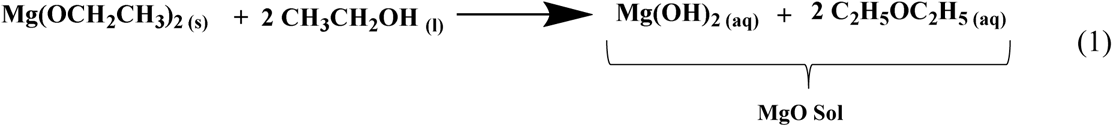

The nanocomposites were synthesized by adding 130 mL AgNPs solution dropwise into 300 mL MgO Sol under continuous stirring. The mixing ensures the dispersion of the nanoparticles within the MgO Sol. The gel formed was then kept undisturbed for a week in a dark room. The latter was then dried in an oven at a temperature of 60 °C.

### 2.5 Determination of the Minimum Inhibitory Concentration (MIC) and the Minimum Bactericide Concentration (MBC)

The evaluation of antimicrobial activity was conducted on multidrug resistant clinical isolates. This method assesses the sensitivity of bacteria and yeasts to different samples according to the microdilution described in ISO 20776-1, 2006 and amended by the Antibiotic Committee of the French Society of Microbiology (CA-SFM / EUCAST 2019). The technique used is that of microdilution in liquid medium of EUCAST 2019 with some modifications [9]. The principle consists of diluting the suspensions of AgNPs in a liquid culture medium and inoculating this medium with microbial strains to be tested. This was accomplished in the following manner:1 mL of the sample was prepared at a concentration of 10 mg/mL and 4 mL of Muller Hinton Broth (MHB) added to the culture medium; A 96-well microplate was used. 100 µL of the previous mixture was distributed in the wells of the first column in triplicates; 50 μL of the medium previously prepared is then distributed in the wells, from the second column to the 12^th^ in triplicates as well; a successive geometric progression dilution of a ratio of two of the sample is carried out. This is from the first well to the tenth well; the 10^th^, 11^th^ and 12^th^ wells consisted of the negative control (sample + culture medium), the positive control (inoculum + culture medium) and the sterility control (culture medium only) respectively; 50 μL of a preparation consisting of 130 μL of inoculum at 0.5 mL Mc Farland and 19.85 mL of culture medium is added to each then preceding; The concentration range of the samples in the reaction medium is 1; 0.5; 0.25; 0.125; 0.0625; 0.0312; 0.0156; 0.0078; 0.0039; 0.0019 (mg / mL). While that of the microorganisms is 5×105 CFU/mL for bacteria; The microplates were incubated at 35 °C for 18 - 24 hours and the MIC was considered the smallest sample concentration for which no growth was observed;

In bacteria, tetrazolium chloride was used as a color indicator. The MBC was determined by transplanting 50 µL of suspension from each well where no shoots were observed in the corresponding 50 µL crop medium. This was the smallest concentration for which no shoots were observed after transplanting. From these reports, the bacteriostatic or bactericidal nature of the various substances was confirmed. When these ratios are ≥ 4, the substance is said to be bacteriostatic; if these ratios are< 4, the substance is bactericidal. If they are =1, then it is said to be absolute bactericidal.

## 3. RESULTS AND DISCUSSION

### 3.1. Synthesis

*Talinum triangulare*leaves were surface cleaned with running tap water followed by deionized water to remove all the dust and unwanted visible particles. The aqueous extract of *T. triangulare* was prepared by boiling 25g of finely cut leaves in 250 mL deionized water for 5 min at 80 °C and stirred for 5 min using a hot plate equipped with a magnetic stirrer [12–14]. The extract was filtered through Whatman No. 1 filter paper and cotton to remove particulate matter.A pale green solution was obtained. This pale green solution later changes to pale brown on cooling. It was stored at 4 °C for further use. This extract is usable for 1 week due to the gradual loss of plant extract viability for prolonged storage. Figure 2 presents a change in color after the addition of silver nitrate to the aqueous extract. The formation of AgNPs was observed from the fifth minute of incubation by a change in coloration from pale brown to pale yellow. This change in coloration is due to the reduction ofAg^+^ ions in AgNO_3_ by secondary metabolites.

**Figure 2:**
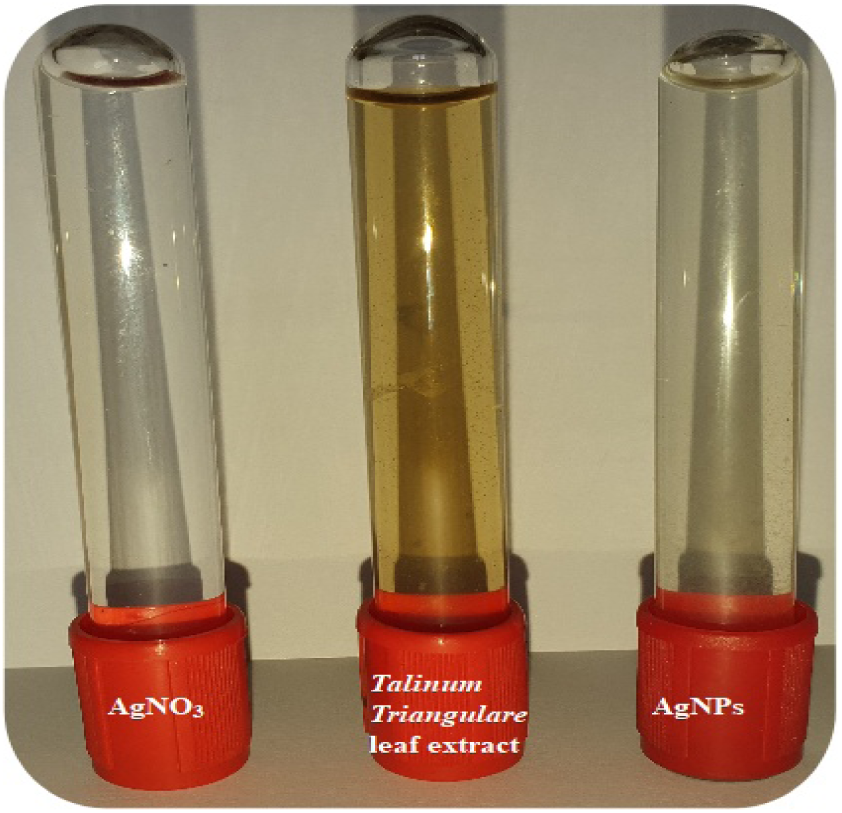
Silver nitrate, *Talinum triangulare* leaf extract and silver nanoparticles solution

A cream white powder was obtained after the synthesized AgNPs solution was mixed with MgO Sol. The mixture was allowed to undergo gelation, where the sol transformed into a gel-like structure. The gel was then aged to allow for further stabilization and maturation of the nanocomposite structure. The latter was dried in an oven at 60 °C to remove any remaining solvent and formation of the final nanocomposite structure.

### 3.1. Ultraviolet Visible Spectroscopy

The formation of nanoparticles was confirmed by the UV-Vis spectroscopy with a SPR band between 350 and 550 nm. This Plasmon resonance band is due to an optical property of the AgNPs related to the exaltation of the electric field, which puts the conducting electrons in uniform motion. The large width of this band is related to a large distribution of the size of nanoparticles [15]. AgNPs formed from the fifth minute of incubation and are formed until 24 hours of incubation period when they reach the plateau and remain stable until 1 week without signs of coalescence (Figure 3). This stability is believed to be due to the formation of ligands that are born or made up of secondary metabolites around the reduced silver atom. The evolution of AgNPs formation was monitored under varying physical conditions such as; time, pH, time, silver nitrate concentration, and extract volume.

**Figure 3:**
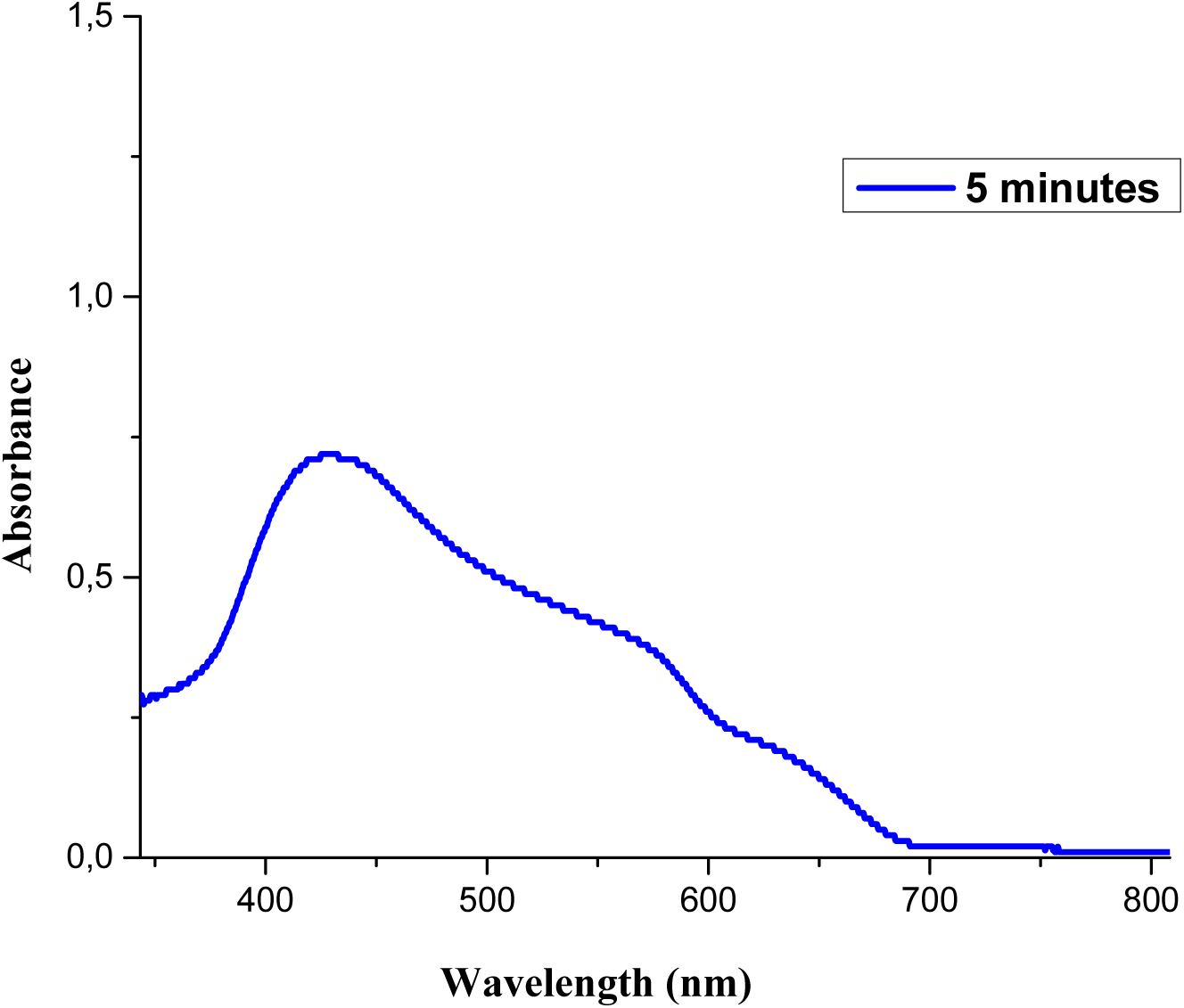
UV-Vis of the AgNPs

#### Influence of pH

The pH level of the reaction medium is crucial in the biosynthesis of green nanoparticles using leaf extracts. In the synthesis of silver nanoparticles (AgNPs), lower pH (acidic and neutral conditions, pH 2, 5, 7) leads to slower reduction rates, larger particle sizes, and reduced stability. Higher pH (alkaline condition, pH 9, 12) promotes faster reduction rates, efficient formation, and stabilization of nanoparticles by preventing aggregation (Figure 4). The alkaline pH also results in smallerand more uniform nanoparticles. Overall, maintaining an optimal pH level, such as pH 12, enhances the biosynthesis of green nanoparticles, ensuring efficient reduction, stability, and control over size and shape.

**Figure 4:**
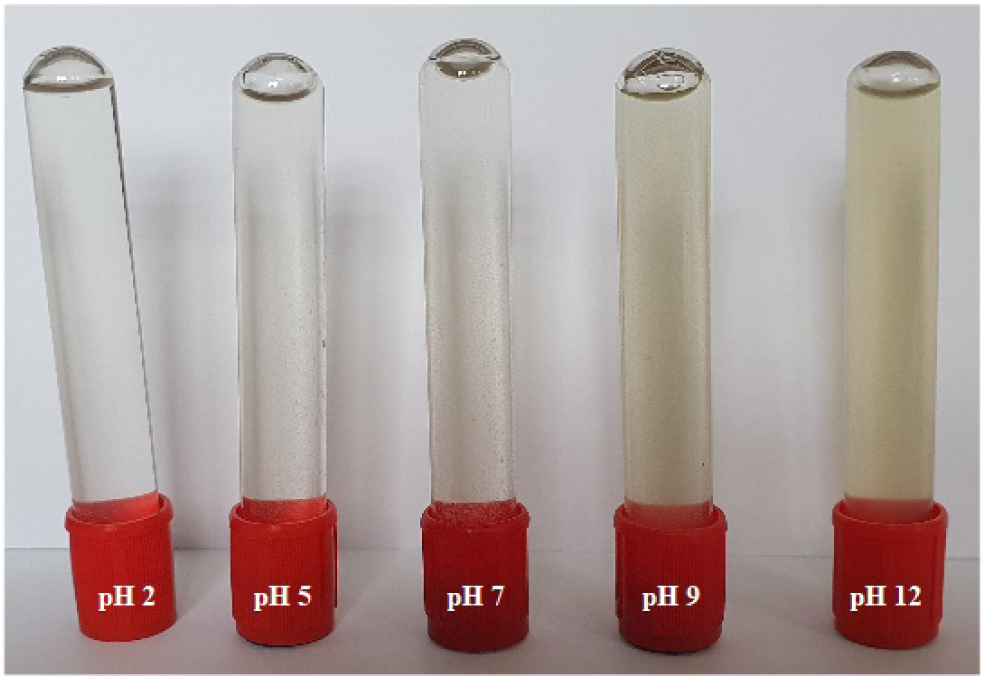
Influence of pH

#### Influence of Time

The time factor is crucial in the biosynthesis of green nanoparticles using leaf extracts and AgNO_3_ precursor, affecting their formation, growth, and stability. Initially, between 5 minutes to 24 hours, silver ions reduce to form primary nuclei growing into nanoparticles, with maximum size and stability at 24 hours (Figure 5). Extending the time to 3 days allows further growth and stabilization but risks agglomeration. At 1 week, there is increased aggregation risk and potential changes in nanoparticle properties. 24 hours is optimal for efficient reduction, nucleation, and growth while maintaining stability and control, though specific applications might require different timings.

**Figure 5:**
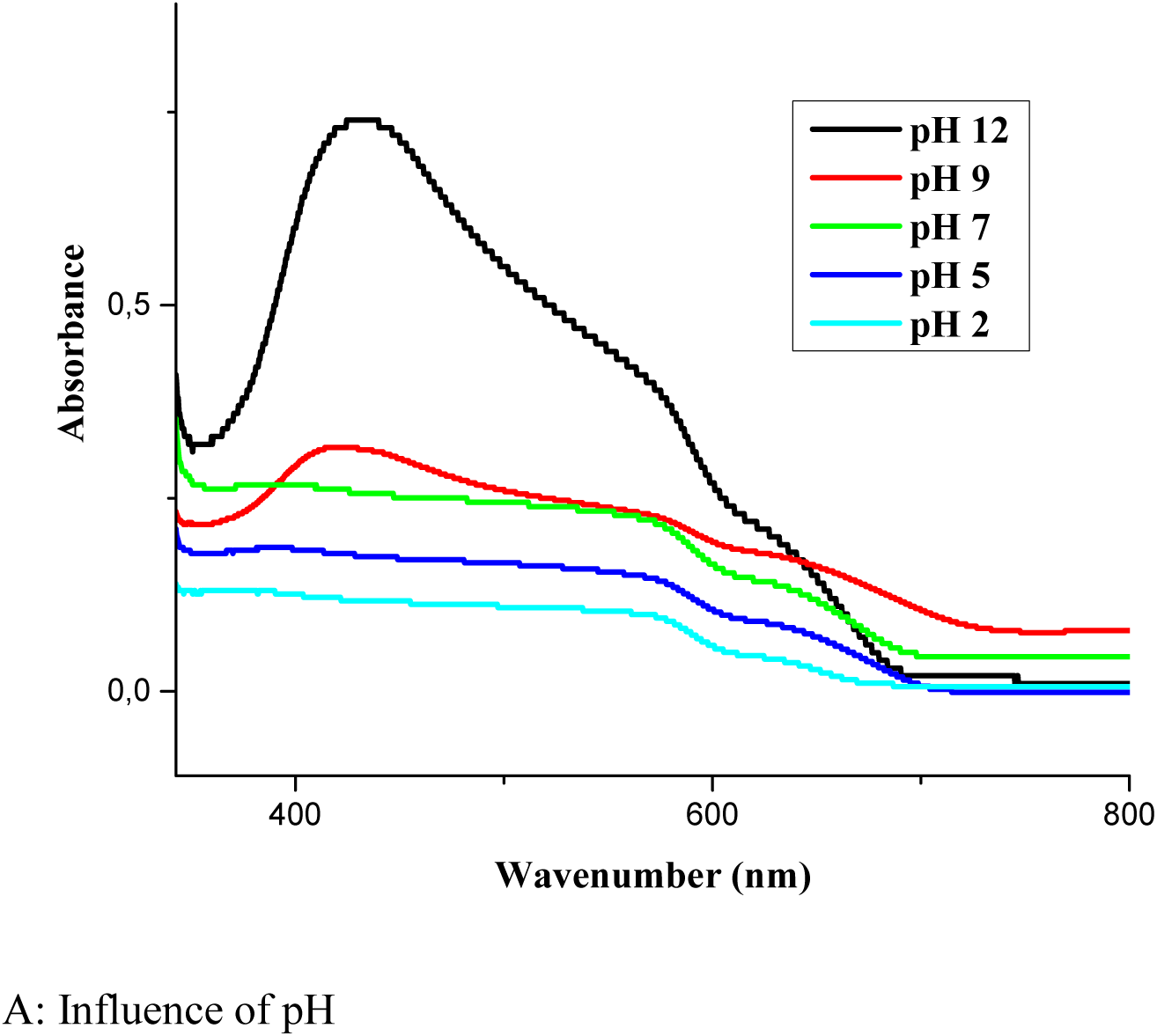
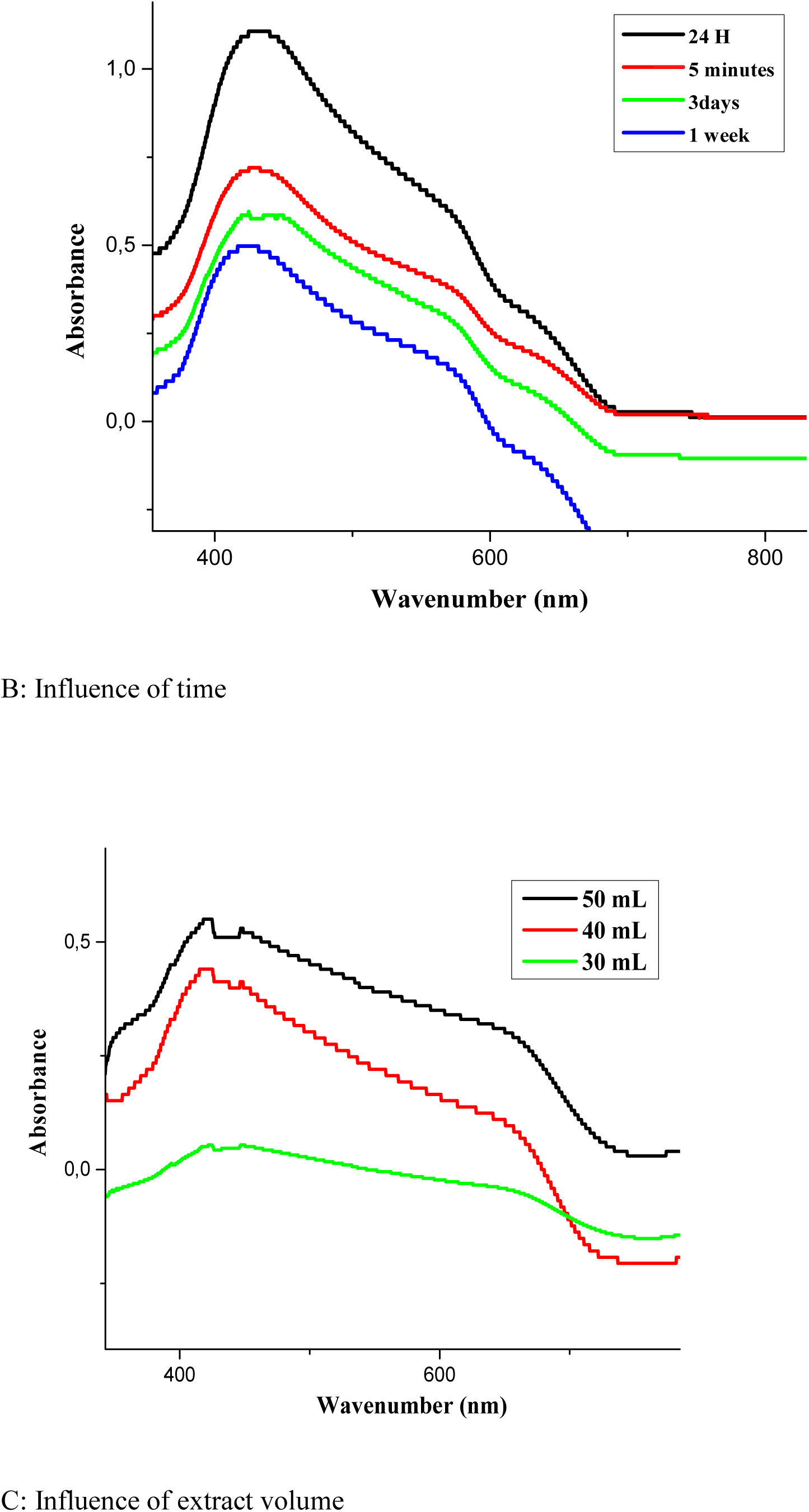
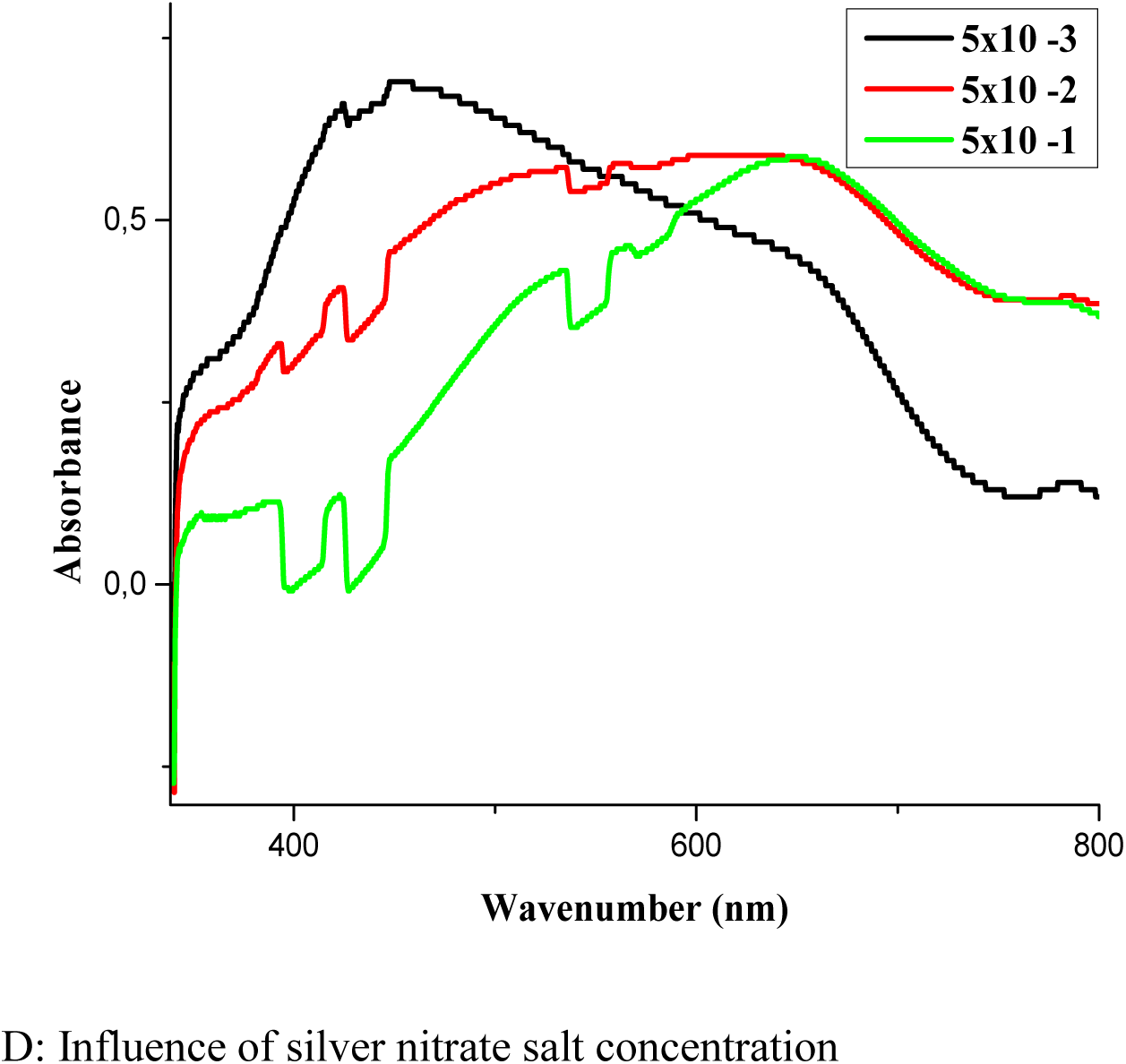
A Influence of pH, B Influence of time, C Influence of extract volume, D Influence of silver nitrate salt concentration

#### Influence of the Extract Volume

The volume of leaf extract in the biosynthesis of green nanoparticles from AgNO_3_affects their formation and properties. Optimal intensity was found at 50 mL of extract. At 30 mL, fewer bioactive compounds lead to slower nanoparticle formation and lower UV-Vis spectrum intensity (Figure 5). Increasing to 40 mL improves nanoparticle formation and intensity but is still less optimal than 50 mL. The 50 mL extract provides an ideal concentration of bioactive compounds and reducing agents, resulting in higher nanoparticle yield, intense absorbance at 428 nm, and desirable size, shape, stability, and uniformity. Adjusting the extract volume is key to optimize the nanoparticle synthesis for specific applications.

#### Influence of Concentration

The concentration of AgNO_3_ precursor significantly influences the biosynthesis of green nanoparticles using leaf extracts (Figure 5). Lower concentration (0.005 M) results in slower nucleation, smaller particle sizes, and broader size distribution, but better stability. Increasing to 0.05 M enhances nucleation and growth, yielding larger nanoparticles with narrower size distribution, though with a higher risk of agglomeration. At 0.5 M, the rapid nucleation and growth produces larger, non-uniform nanoparticles, with increased aggregation and reduced stability due to overcrowding. Optimal concentration balances efficient reduction, growth, and nanoparticle stability. Finally, the optimal synthesis conditions for the formation of stable AgNPs are; pH12, AgNO_3_ concentration of 5×10^-3^ mol/L and a ratio volume of 5:10 between the aqueous extract and AgNO_3_respectively.

### 3.1. X-ray diffraction

The X-ray pattern of AgNPs revealed a series of crystalline phases at (2Ɵ) 08.7°, 18.5°, 21.0°, 24.7°, 27.3°, 34.1°, 38.1°, 43.0°, 44.4°, 51.0°, 58.6°, 61.9°, 64.4°, 71.6°, 81.3°, and 114.0° (Figure 6). The average crystalline of the synthesized nanoparticles was determined using the Debye-Scherrer equation 2;

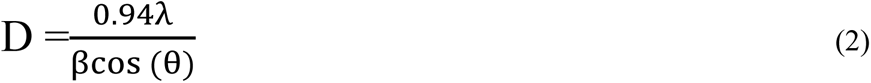

**Figure 6:**
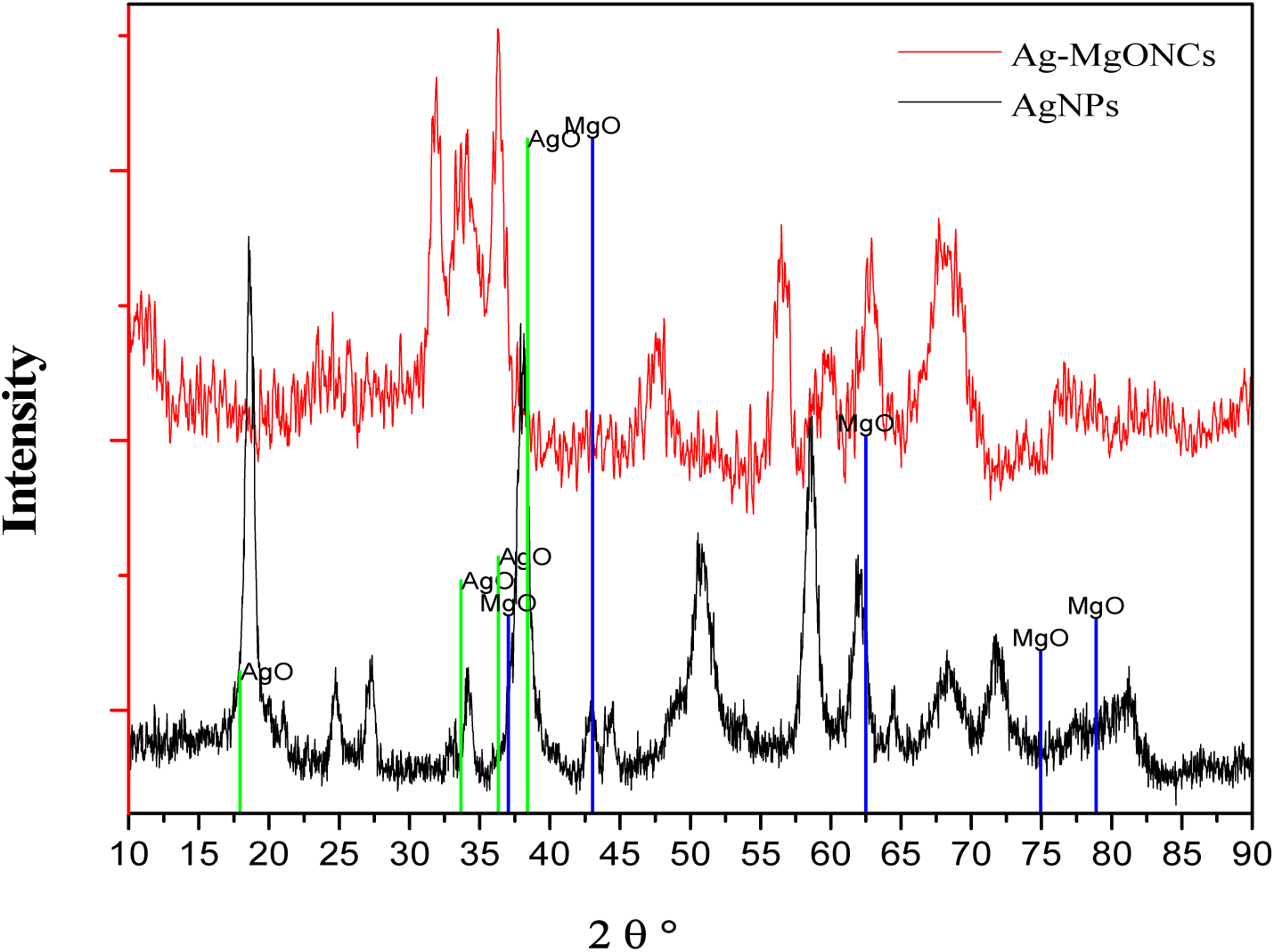
XRD Spectrum of AgNPs/Ag-MgONCs.

Where D is the crystalline size (nm); λ is the X-ray wavelength (λ = 1.5406 Å); β is the Full Width at Half Maximum (FWHM) of the diffraction peaks measured in radians; θ is the Bragg diffraction angle. No other characteristic peaks were found in the XRD spectra, indicating the high purity of the prepared Ag@Ag_2_O nanoparticles (Figure 6).To calculate the average crystalline particle size of the synthesized Ag@Ag_2_O nanoparticles, we preferred intense peaks of Ag and Ag_2_O. 2θ value of 38.1° that can be indexed to the (0 1 1) plane of the face-centered cubic (FCC) structure (JCPDS file: 65-2871) [13]confirming the hexagonal crystalline states of the nanoparticles and giving an average particle size of 37.1 nm.

On the other hand, the X-ray pattern of Ag-MgONCs revealed a series of crystalline phases at (2Ɵ) 08.6°, 27.5°, 32.2°, 38.1°, 44.3°, 46.2°, 64.4°, 77.2°, and 114.0°.The PXRD pattern showed the presence of two phases, the cubic phase of MgO at 2θ value; 38.1°, 43,0°, 62,4° and 77.2° corresponding to the (111, 200, 220) and (222) planes, respectively (JCPDS file: 31-12380) confirming the cubic crystalline states of the nanocomposites. It also showed the presence of cubic phase of Ag_2_O at 2θ values;8.6°, 27.5°, 32.2°, 46.2°, and 114.0° corresponding to the (200, 110, 111, 220) and (133) planes, respectively (JCPDS file: 31-12380) confirming the cubic crystalline states of the nanocomposites.The calculatedcrystalline particle size gave 9.1 nm.

### 3.1. Micrographic Images and Microanalysis of AgNPs and Ag-MgONCs

Figure 7 shows the surface morphology of green synthesized AgNPs. The images give a spherical shape for the entire sample with a cluster of aggregation. The aggregation reduces gradually as the concentrations increases. The EDX, as shown in Figure 7, shows all the elements present in the compound. The elemental analysis confirmed the reactant composition of all elements, with Ag having the broadest peak at energy 3.7 Kev. The SEM-EDX determines the elemental compositions of AgNPs, (Figure 7).

**Figure 7:**
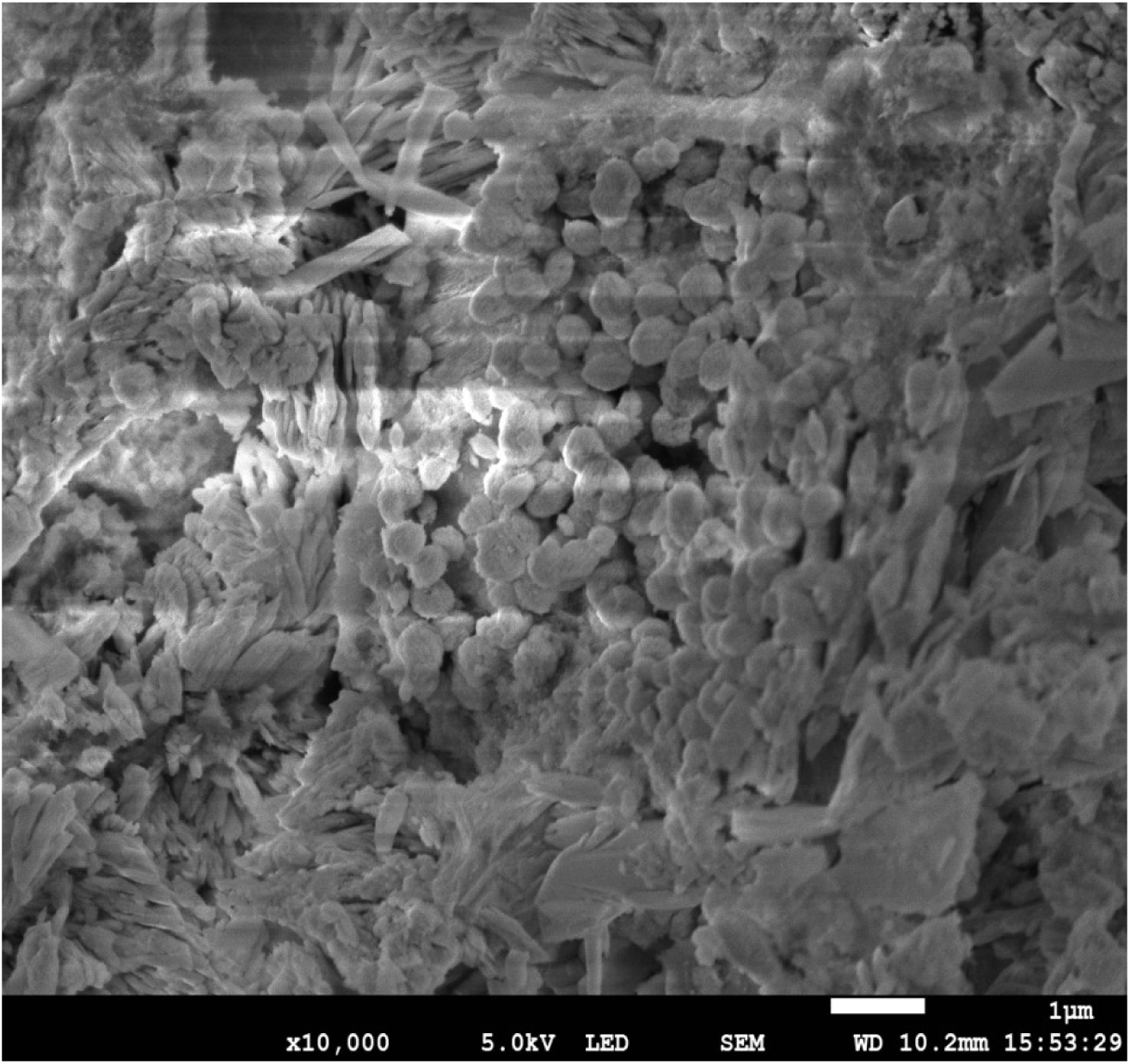
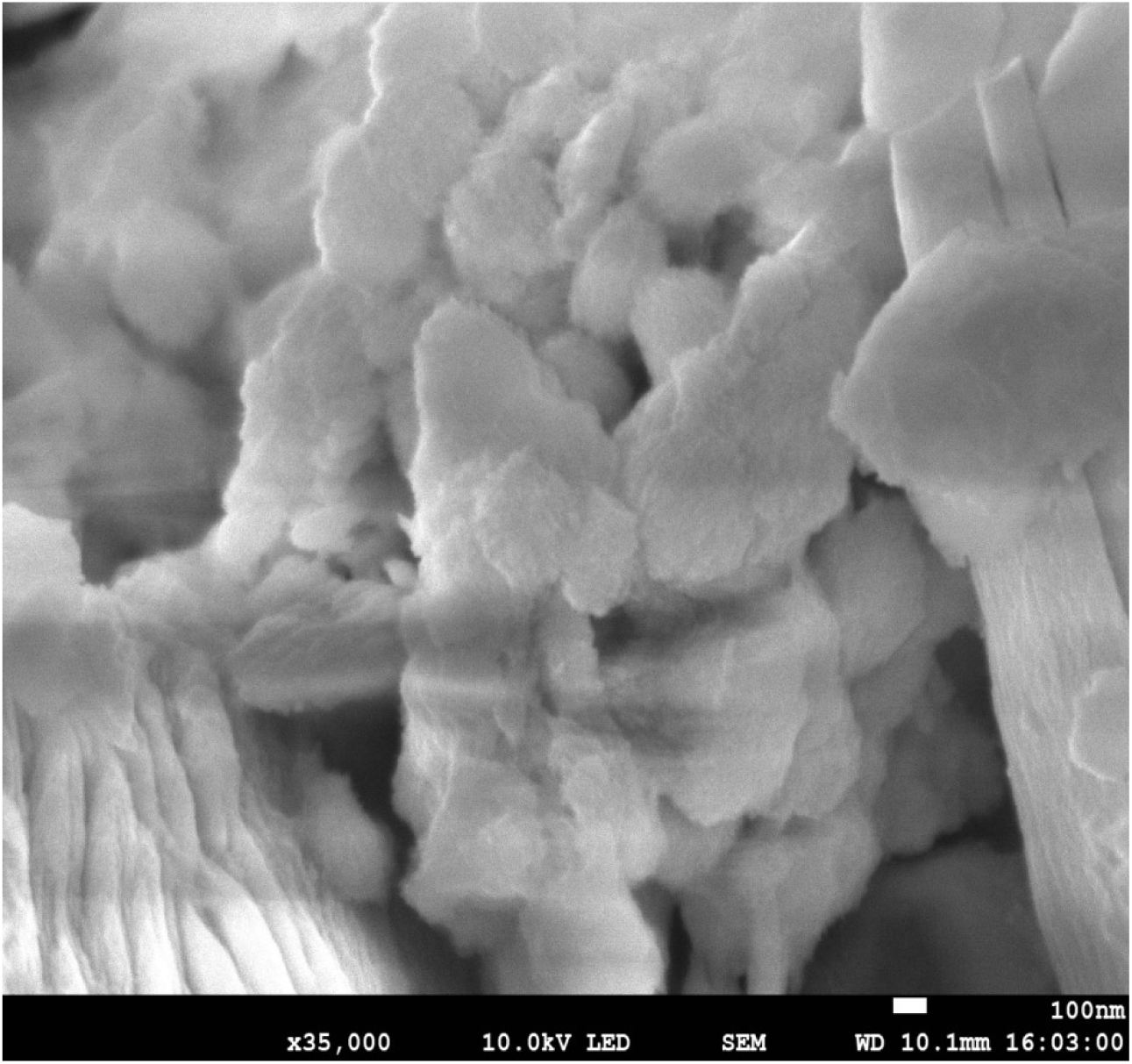
Micrographic images of AgNPs x10000 and x35000

The pattern of the grain with several sharp rings confirmed that the AgNPs nanostructure is highly polycrystalline in nature. Carbon, oxygen, and nitrogen have been elucidated in AgNPs. These are the main atoms that make up organic matter, so they match the composition of most of the metabolites identified in the plant extract used. In addition, the presence of carbon and oxygen testifies to the stabilization of AgNPs by organic molecules (secondary metabolites). These results agree with those of Belle EbandaKediet al. in 2018 on silver nanoparticles [13].

SEM of theAgNPs consisted of subjecting them to a high-energy electron beam and detecting the backscattered electrons.Figures 7 shows the images of AgNPs at a resolution of 100 nm. This result is an aggregate of particles of spherical shapes whose reflection of light shows the crystalline nature of the powder. Energy dispersive x-ray spectroscopy highlights the atomic composition of these AgNPs as shown in Table 1.

**Table 1:** Atomic % Composition of AgNPs.

| Element | Wt.% | Error |
| --- | --- | --- |
| C | 16.59 | ± 1.31 |
| Mg | 18.12 | ± 3.42 |
| O | 59.82 | ± 3.14 |
| Ag | 5.46 | ± 1.99 |
| Pd | 0.01 | ± 1.12 |
| Total | 100.00 |  |

Scanning microscopic analysis of the nanoparticles revealed a spherical aggregate of AgNPs. This is consistent with the work of several researchers in this field like Eya’aneMeva et al. in 2017 [12]. Energy dispersion X-ray spectroscopy has revealed the different atoms present in the silver nanoparticles, depending on the database of the device used. Carbon, oxygen, and magnesium have been elucidated in AgNPs. These are the main atoms that make up organic matter, so they match the composition of most of the secondary metabolites identified in the plant extract used. In addition, the presence of carbon and oxygen testifies to the stabilization of AgNPs by organic molecules (secondary metabolites).

Figure 9 shows the SEM images of Ag-MgONCs at a resolution of 1µm. Energy dispersive X-ray spectroscopy and element mapping highlights the atomic composition of these Ag-MgONCs (Figure 10; Table 2). Energy dispersive X-ray spectroscopy revealed the atoms oxygen and magnesium at the majority. These are the main components of the nanocomposite matrix.

**Figure 8.**
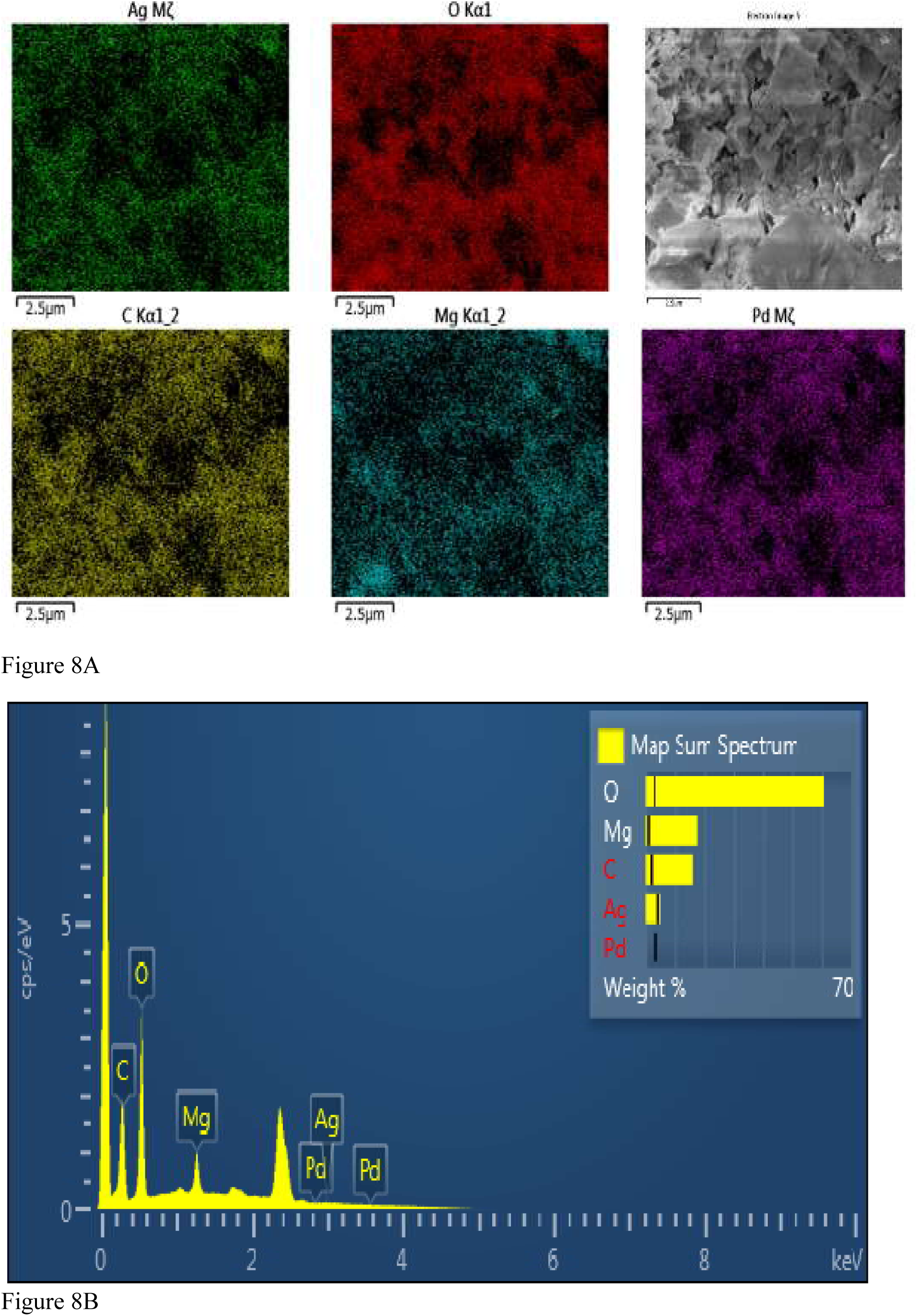
A, B: EDX spectrum of AgNPs

**Figure 9:**
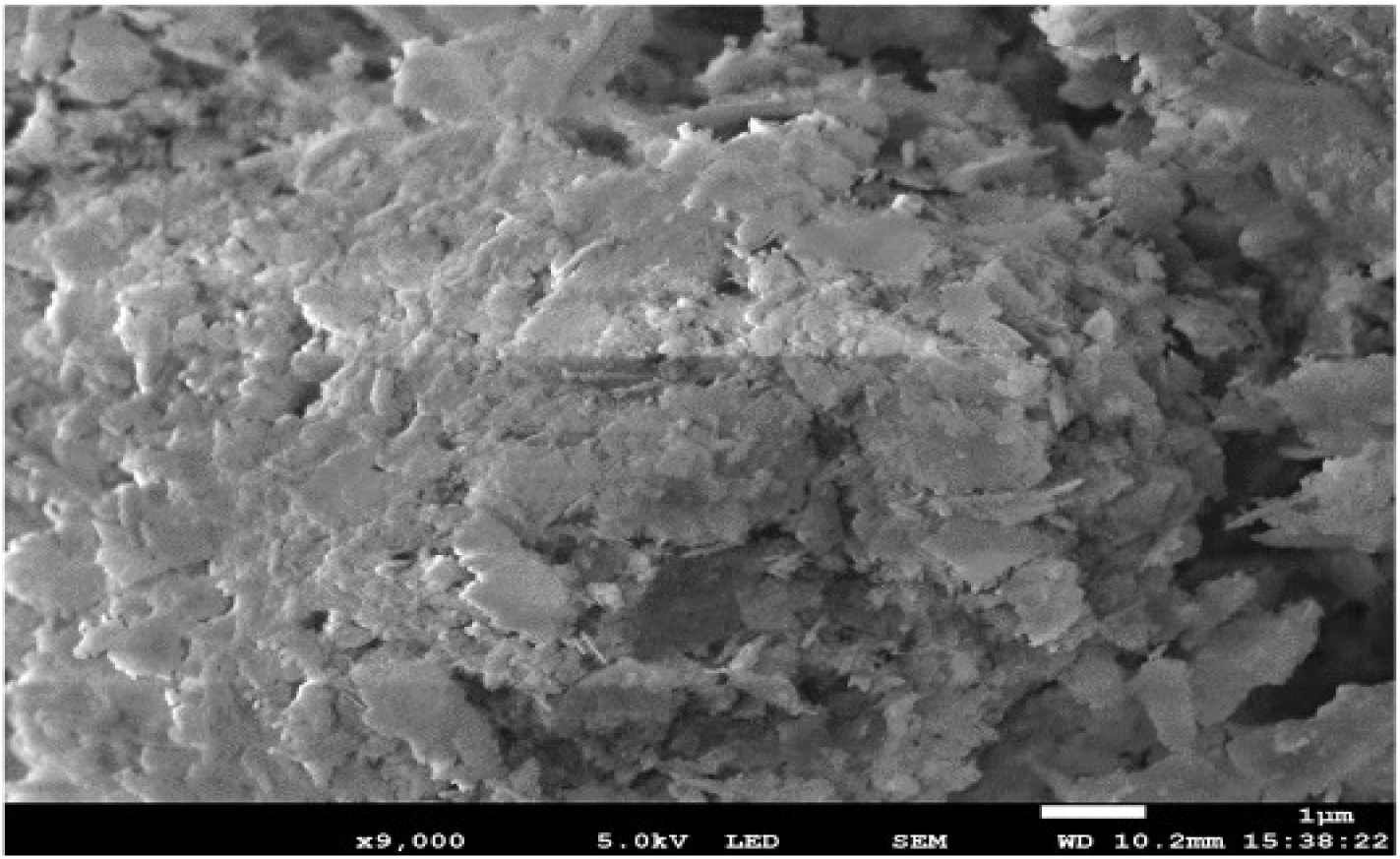
Micrographic images of Ag-MgONCs

**Figure 10:**
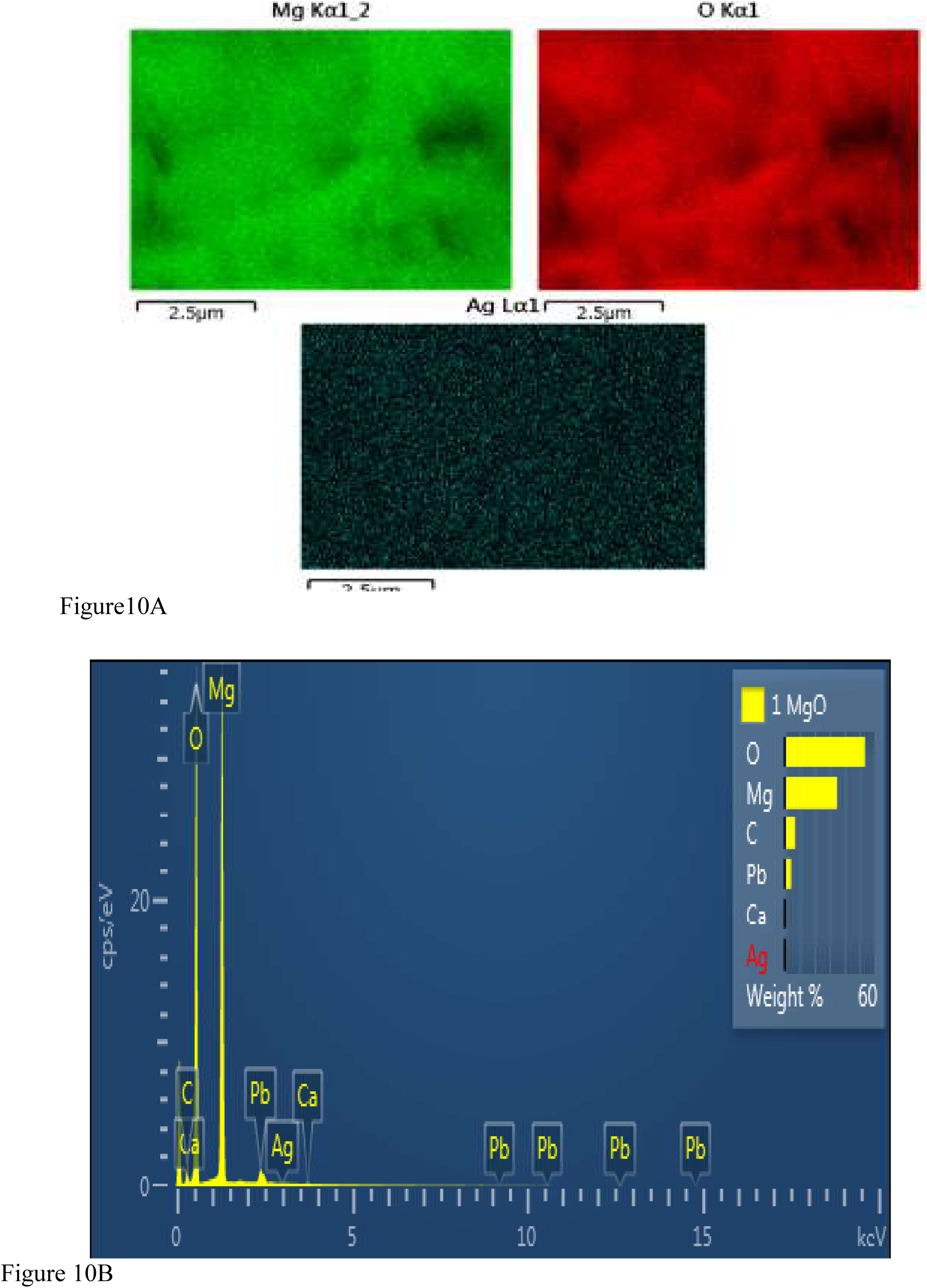
A, B: EDX spectrum of Ag-MgONCs

**Table 2:**
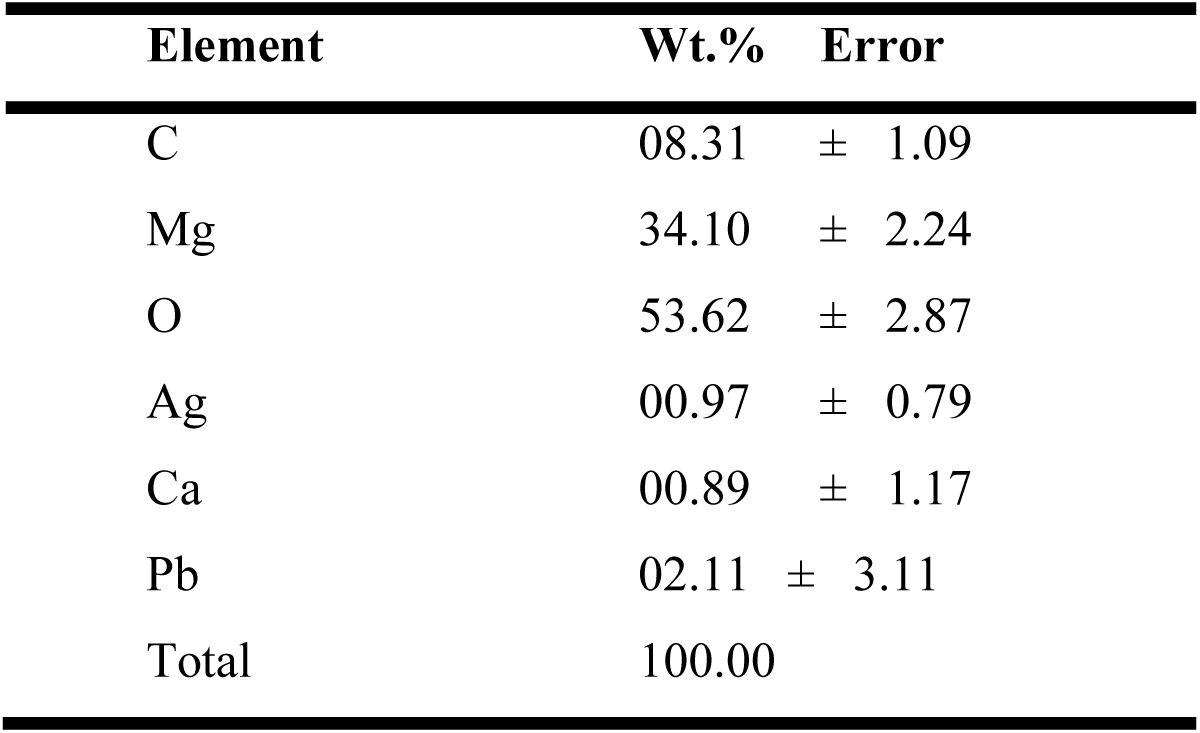
Atomic % Composition of Ag-MgONCs.

### 3.2. Dynamic light scattering analysis of AgNPs and Ag-MgONCs

The nanomaterials dispersed in a colloidal solution are in continuous Brownian motion. DLS measures the light scattering as a function of time, which combined with the Stokes-Einstein assumption, is used to determine the nanoparticle hydrodynamic diameter (i.e., diameter of the nanomaterial and the solvent molecules that diffuse at the same rate as the colloid) in solution. Figure 11 shows size distribution by Intensity graphs that elucidate the hydrodynamic diameter of the different nanomaterials.

**Figure 11:**
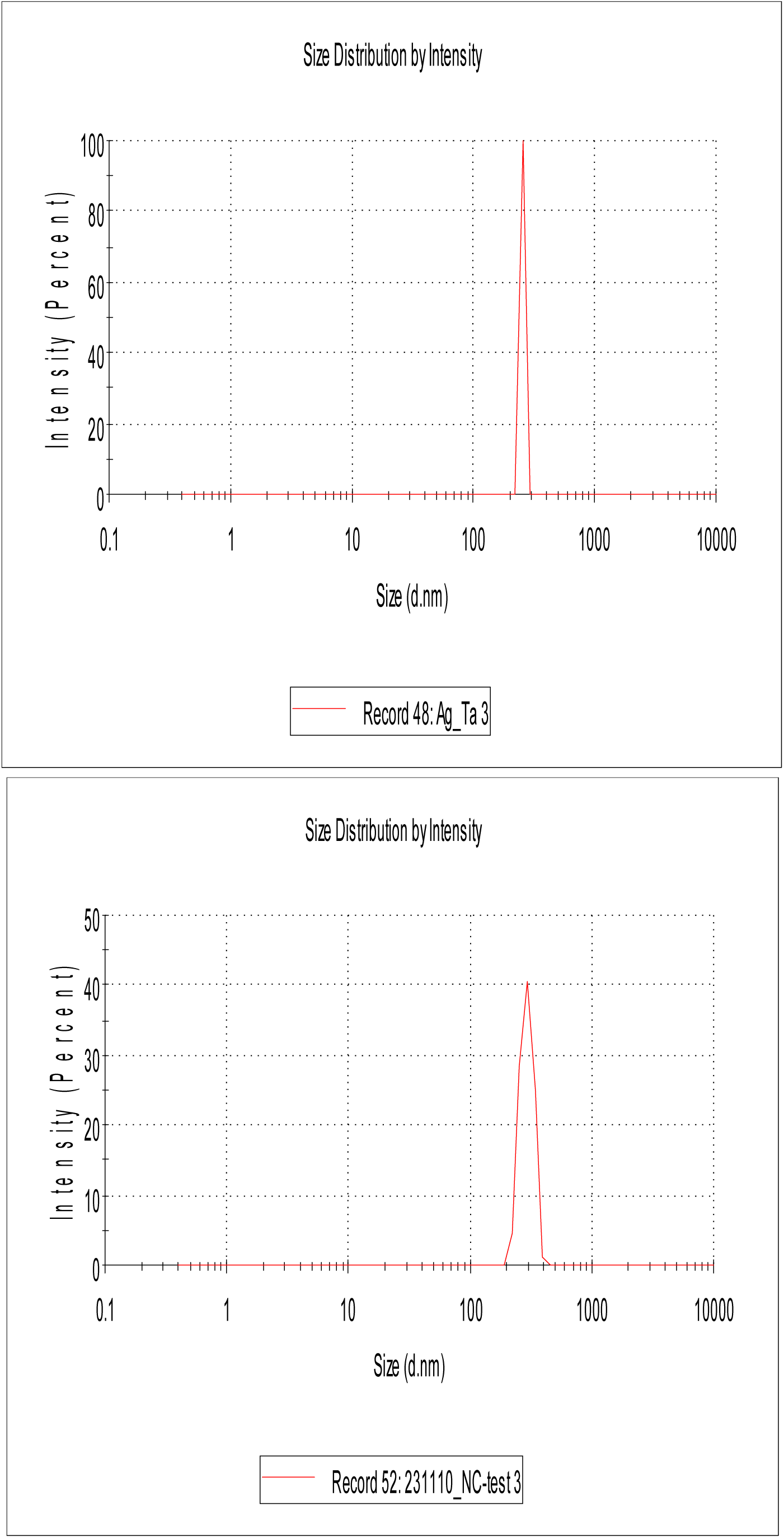
Size Distribution by Intensity of AgNPs(A) and Ag-MgONCs (B)

The graphs demonstrated agglomerate sizes of 295 nm and 225 nm for Ag-MgONCs and AgNPs, respectively. Given the respective sizes of the nanomaterials with radius within the range 100 nm – 300 nm we could conclude that we have stable solutions of aggregated nanomaterialswith a polydispersity index of approximately 0.3.

### 3.3. Microbiological Assays

#### Solubility test

A solubility test was carried out (Table 3). Dimethyl Sulfoxide, Distilled water, Ethanol, and Tween (C_26_H_50_O_10_) were used. On analysis, distilled water had the best dissolution properties.

**Table 3:** Solubility test.

| Samples | Distilled H<sub>2</sub>O | DMSO | Ethanol | Tween (C<sub>26</sub>H<sub>50</sub>O<sub>10</sub>) |
| --- | --- | --- | --- | --- |
| Ag-MgONCs | + | + | - | - |
| AgNPs | + | - | - | - |
| Aqueous Extract | + | - | - | + |

Not soluble (-) ; Soluble (+)

The resistance of human pathogens to commercially available antimicrobial agents and antibiotics has raised the need to explore new natural and inorganic substitutes to overcome the problem [16]. Anh-TuanLe *et al*. [17] demonstrated that Ag nanomaterials were attached to the cell surface of the bacteria strains and then penetrated the cell, destroyed the cell cytoplasm, and killed the organism. They also found that Ag nanomaterials significantly increase cell permeability and affect the proper transport through the plasma membrane. Sulfur and phosphoruscontaining proteins or enzymes or phosphorus moiety of DNA of bacterial system may be affected by Ag nanomaterials, leading to the inhibition of enzyme system of the organism [18].

#### Microbiology control sample

Ag-MgONCs, AgNPs and the aqueous extract have been subjected to microbiological control to ensure that they are not contaminated with germs that may grow on culture media used for antimicrobial testing. Table 4 represents observations of culture media impregnated with samples after 48 hours of incubation. After seeding the samples inMuller Hinton and MBE media, no shoots were observed after 48 hours of incubation.

**Table 4:** Microbiological control of samples.

| Samples | Muller Hinton | MBE |
| --- | --- | --- |
| Ag-MgONCs | A | A |
| AgNPs | A | A |
| AqueousExtract | A | A |

A : Lack of shoots

#### The minimum inhibitory and bactericidal concentrations

The evaluation of the antimicrobial activity of the synthesized sample of the different bacteria strains gave values of MIC and MBC, which we compared with the reference (Ciprofloxacin and Gentamycin). These values are recorded in Tables 5 and 6, respectively.

**Table 5:** MIC of the sample tested on the strains compared to Ciprofloxacin and Gentamycin.

| MIC (mg/mL) |  |  |  |  |  |
| --- | --- | --- | --- | --- | --- |
| Bacteria strains<br>Test milieu | *Escherichia coli* | *Acinetobacter baumannii* | *Pseudomonas aeruginosa* | *Klebsiella pneumoniae* | *Proteus mirabilis* |
| Aqueous Extract | #1.0000 ± 0.0000 | #1.0000 ± 0.0000 | #1.0000 ± 0.0000 | #1.0000 ± 0.0000 | #1.0000 ± 0.0000 |
| AgNPs | #0.5000 ± 0.0000 | #1.0000 ± 0.0000 | #1.0000 ± 0.0000 | #1.0000 ± 0.0000 | #0.5000 ± 0.0000 |
| Ag-MgONCs | #0.5000 ± 0.0000 | #1.0000 ± 0.0000 | #1.0000 ± 0.0000 | #1.0000 ± 0.0000 | #0.2500 ± 0.0000 |
| Cipro | 0.5000 ± 0.0000 | 0.0156 ± 0.0000 | 0.1250 ± 0.0000 | 0.0156 ± 0.0000 | 0.0312 ± 0.0000 |
| Gentamycin | 0.5000 ± 0.0000 | 1.0000 ± 0.0000 | 1.0000 ± 0.0000 | 0.0156 ± 0.0000 | 0.0625 ± 0.0000 |
| Positive control | - | - | - | - | - |
| Negative control | + | + | + | + | + |
| Sterility control medium | - | - | - | - | - |

No shoot (-); The presence of a shoot (+); Bactericidal (#)

**Table 6:**
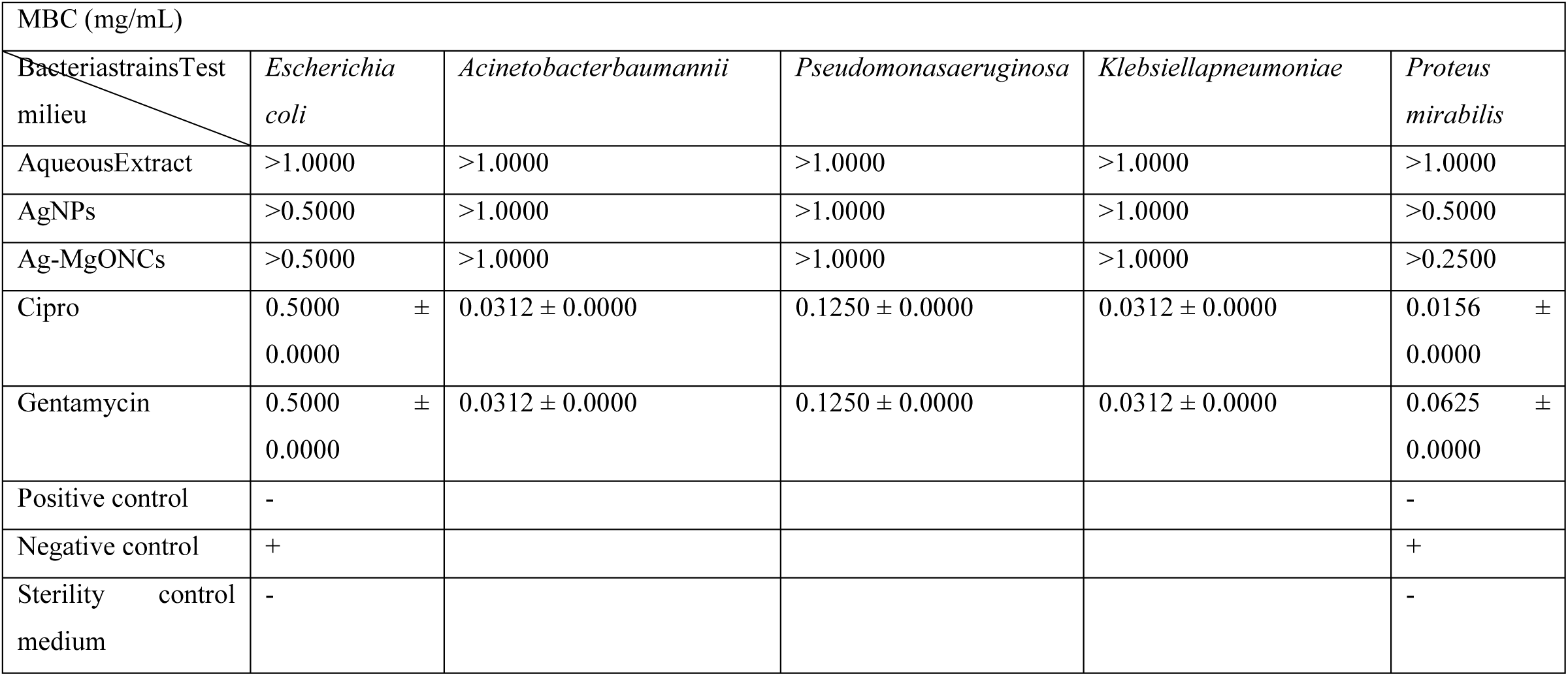
MBC of the sample tested on the strains compared to Ciprofloxacin and Gentamycin.

#### Aqueous Extract Analysis

The extract showed little antimicrobial activity against the multidrug-resistant strains (Table 6). The evaluation of the MBC of the synthesized AgNPs shows that they have low bactericidal properties. The determination of the MIC/MBC of the aqueous extract was carried out in comparison with that of the aqueous extract, Ag-MgONCs, and reference drugs.

#### AgNPs analysis

AgNPs appear to have the same MIC against *E. coli*, and a measurable MIC against the other strains compared to those of the reference drugs (Figure 12). The evaluation of the MBC of the synthesized AgNPs shows that they are bactericidal against multiresistant *E. coli*.

**Figure 12:**
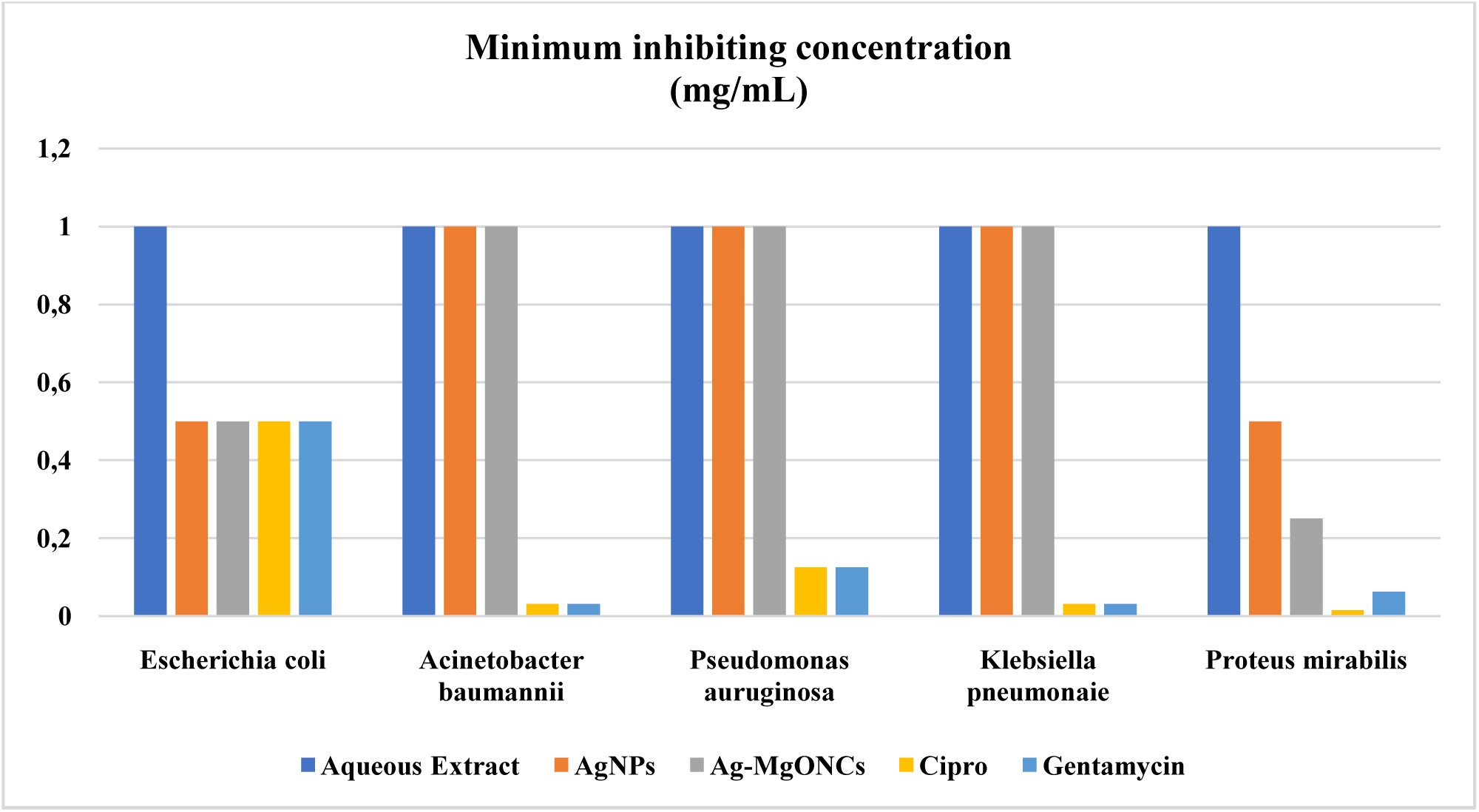
Minimum inhibiting concentration (mg/mL)

The determination of the MIC/MBC of AgNPs was carried out in comparison with that of the aqueous extract, Ag-MgONCs, and reference drugs. These nanoelements demonstrated interesting antibacterial activities with respect to the multiresistant strains (*Escherichia coli, Acinetobacter baumannii, Pseudomonas aeruginosa, Klebsiella pneumoniae and Proteus mirabilis).*The nanoparticles however, showed low activity for *Acinetobacter baumannii,Pseudomonas aeruginosa, Klebsiella pneumoniae, and Proteus mirabilis* bacterial strains compared to those of the references. These activities are related to the possibility of releasing the silver ions directly into the microbial cell, which will be able to act on various sites. Synthesized nanoparticles are covered with extract molecules that may also have intrinsic antimicrobial properties [19]. Therefore, AgNPs would also play a good role as a vector for delivering Ag^+^ ions and molecules from the extract at their site of action.

#### Ag-MgONCsAnalysis

Allegedly, Ag-MgONCs have the matching MIC against *E. coli* and *P. mirabilis*, and a striking MIC against the other strains compared to those of the reference drugs (Figure 12). Pronounced activity with *E. coli* and *P. mirabilis* against the nanocomposites was perceived compared to the aqueous extract and AgNPs. We observed a minimum inhibition concentration (0.500 mg/mL) and (0.250 mg/mL) the same for *E. coli* and higher for *P. mirabilis* compared to those ofreference molecules (0.500 mg/mL) and (0.0312 mg/mL) respectively. In view of its minimum bactericide concentration, it is bactericidal.This confirms our hypothesis of the latter having improved properties. Nevertheless, the reference drugs are one-step ahead.

The determination of the MIC/MBC of Ag-MgONCs was carried out in comparison with that of the aqueous extract, AgNPs, and reference drugs. These nanoelements demonstrated interesting antibacterial activities with respect to the multiresistant strains (*Escherichia coli, Acinetobacter baumannii, Pseudomonas aeruginosa, Klebsiella pneumoniae and Proteus mirabilis).*However, the nanocomposites showed low activity for *Acinetobacter baumannii, Pseudomonas aeruginosa, and Klebsiella pneumoniae* bacterial strains compared to that of the references. These activities are related to the possibility of releasing the silver ions directly into the microbial cell, which will be able to act at various sites. Synthesized nanomaterials are covered with molecules from the extract that may also have intrinsic antimicrobial properties [19]. Ag-MgONCs therefore would also play a good role as a vector for delivering Ag^+^ ions and molecules from the extract at their site of action.

## 4. CONCLUSION

Ag-MgO nanocomposites mediated from *Talinum triangulare* leaf extract have been designed. The utilization of PXRD and DLS techniques enabled the determination of particle sizes consistent with the dimensional criteria for nanomaterials. SEM-EDX analysis provided spherical morphology for the nanoparticles and granular morphology for the nanocomposites. Collectively, these methods afforded comprehensive insights into the synthesis of nanocomposites. The synthesized nanocomposites demonstrated bactericidal properties, with measurable minimum bactericidal concentration values comparable to those of reference medicines. Additionally, the nanocomposites exhibited enhanced properties relative to their bulk counterparts while maintaining eco-friendly characteristics. Future endeavors include conducting TEM analysis to further validate our findings and exploring in vivo studies on live albino rats as a subsequent application.

## Declarations

### Competing Interests

Authors are required to disclose financial or non-financial interests that are directly or indirectly related to the work submitted for publication.

Data availability The authors declare that the data supporting the findings of this study including raw data files are available from the corresponding author upon reasonable request.

All authors contributed to the study conception and design. Material preparation, data collection and analysis were performed by Fonye Nyuyfoni Gildas, Paboudam Gbambie Awawou, Kwati Leonard, Gilbert Njowir Ndzeidze. The first draft of the manuscript was written by Fonye Nyuyfoni Gildas, Paboudam Gbambie Awawou and Eya’ane Meva Francois, all authors commented on previous versions of the manuscript. All authors read and approved the final manuscript.

### Conceptualization

Paboudam Gbambie Awawou, Eya’ane Meva Francois Investigation: Fonye Nyuyfoni Gildas, Kwati Leonard, Gilbert Njowir Ndzeidze, Methodology: Eya’ane Meva Francois, Paboudam Gbambie Awawou Resources: Eya’ane Meva Francois, Paboudam Gbambie Awawou

## REFERENCES

[1] F Eya’ane Meva, C Okalla Ebongue, S V Fannang, M L Segnou, A A Ntoumba, P Belle Ebanda Kedi, R-E Njike Loudang, A Yonga Wanlao, E R Mang, E A Mpondo Mpondo. Natural Substances for the Synthesis of Silver Nanoparticles against Escherichia coli: The Case of *Megaphrynium macrostachyum* (Marantaceae), *Corchorus olitorus* (Tiliaceae), *Ricinodendron heudelotii* (Euphorbiaceae), *Gnetum bucholzianum* (Gnetaceae), and *Ipomoeabatatas* (Convolvulaceae). Journal of Nanomaterials.2017;(1):1–6. 10.1155/2017/6834726.

[2] W B Ayinde, M WGitary. M Muchindu, A Samie. Biosynthesis of Ultrasonically Modified Ag-MgO Nanocomposite and Its Potential for Antimicrobial Activity. Journal of Nanotechnology 2018;(1):1–10. 10.1155/2018/9537454.

[3] Y H Leung, A M C Ng, X Xu, Z Shen, L A Gethings, M T Wong, C M N Chan, M Y Guo, Y H Ng, A B Djurisic, P K H Lee, W K Chan, L H Yu, D L Phillips, A P Y Ma, F C C Leung. Mechanismsofantibacterial activity of MgO: non-ROS mediated toxicity of MgO nanoparticles towards *Escherichia coli*. Small, 2014;(10):1171–1183. 10.1002/smll.201302434.

[4] X Zhu, D Wu, W Wang, F Tan, P K Wong, X Wang, X Qiu, X Qiao. Highly effective antibacterial activity and synergistic effect of Ag-MgO nanocomposite against *Escherichiacoli*. Journal of Alloys and Compounds, 2016;(684):282–290.10.1007/s12010-014-0852-z.

[5] O VKharissova, HR Dias, BI Kharisov, B OPerez, V M J Perez. The greener synthesis of nanoparticles. Trends in Biotechnology, 2013;(31):240–248. 10.1007/s11356-023-27437-9.

[6] M Shah, D Fawcett, S Sharma, S K Tripathy, GEJ Poinern. Green synthesis of metallic nanoparticles via biological entities. Materials. 2015;(8):7278–7308. 10.3390/ma8115377

[7] L Arangasamy, V Munusamy.Tapping the unexploited plant resources for the synthesis of silver nanoparticles. African Journal of Biotechnology. 2008;(7):3162– 3165. 10.4314/ajb.v7i11.59252.

[8] EE Elemike, D C Onwudiwe, O E Fayemi, A C Ekennia, E Ebenso, L R Tiedt. Biosynthesis, electrochemical, antimicrobial and antioxidant studies of silver nanoparticles mediated by *Talinum triangulare* aqueous leaf extract. J Clust Sci. 2017;(28):309– 330. 10.1007/s10876-016-1087-7.

[9] A O Adewunmi, E A Sofowora. Preliminary screening of some plant extracts for molluscicidal activity. Planta Med. 1980;(39):57–82. 10.1055/s-2008-1074903.

[10] P M Aja, A N C Okaka, P N Onu, U Ibiam, A J Urako.Phytochemical composition of *Talinumtriangulare* (water leaf) leaves. Pak. J. Nutr. 2010;(9):527– 530. 10.3923/pjn.2010.527.530.

[11] M Sigamoney, S Shaik, P Govender, S B N Krishna. African leafy vegetables as bio-factories for silver nanoparticles: a case study on *Amanranthusdubius* c mart. Ex Thell. S. Afr. J. Botany 2016;(103): 230–240. 10.1016/j.sajb.2015.08.022.

[12] F Eya’ane Meva, M L Segnou, M-A Etoh, C. Okalla Ebongue, M H JNko’o, A A Ntoumba, P Belle Ebanda Kedi, R-E, Njike Loudang, S Djiopang Yadou, F A Essombe Malolo, L Ngah, A. Yonga Wanlao, R E Mang, E A Mpondo Mpondo. Simple disclosure of plasmon resonance band maxima by ultraviolet spectroscopy coupled to centrifugation, determination of percentage recovery: a case study of *Gnetum bucholzianum* Engl. leaf mediated silver nanoparticles. Asian journal of biochemical and pharmaceutical research Issue 1, 2017;(7):62–74. 10.24214/AJBPR/7/1/6274.

[13] P Belle Ebanda Kedi, F Eya’ane Meva, L Kotsedi, E L Nguemfo, C B Zangueu, A A Ntoumba, H E Mohamed, A B Dongmo, M Maaza. Eco-friendly synthesis, characterization, in vitro and in vivo anti-inflammatory activity of silver nanoparticle-mediated *Selaginellamyosurus* aqueous extract. International journal of nanomedicine. 2018;(13):8537. 10.2147/IJNAN.S125456.

[14] Z Meiling, D MingLiang, Z Han, X CongSheng. F YaQuin. Synthesis of Silver Nanoparticles in Electrospun Polyacrylonitrile Nanofibers Using Tea Polyphenols as the Reductant. Polymer Engeneering and Sciences. 2013;(53):1099– 1108. 10.1007/s11051-008-9513-x.

[15] N Alam, D Sreeparna, B Shaikh, R Nayan, C Anirban, M Debabrata, N A Begum. Murraya koenegii Spreng. Leaf Extract: An Efficient Green Multifunctional Agent for the Controlled Synthesis of Au Nanoparticles. ACS Sustainable Chemistry and Engeneering. 2014;(4):652–664. 10.1021/sc400562w.

[16] J Annamalai, T Nallamuthu.Green synthesis of silver nanoparticles: characterization and determination of antibacterial potency. Applied Nanoscience. 2016;(6):259– 265. 10.1007/s13204-015-0426-6.

[17] A-T Le, T Tam Le, V Q Nguyen, H Hoang Tran, D A Dang, Q H Tran, D LahmVu. Powerful colloidal silver nanoparticles for the prevention of gastrointestinal bacterial infection. Advances in Natural Sciences: Nanoscience and Nanotechnology, 2012;(3), ArticleID045007. 10.1088/2043-6262/3/4/045007.

[18] G M Sulaiman, W H Mohammed, T R Marzoog, AAA Al-Amiery, AAH Kadhum, AB Mohamad. Green synthesis, antimicrobial and cytotoxic effects of silver nanoparticlesusing*Eucalyptus chapmaniana* leaves extract. Asian Pacific Journal of Tropical Biomedicine. 2013;(3):58–63. 10.1016/S2221-1691(13)60024-6.

[19] R Y Pelgrift, A J Friedman. Nanotechnology as a therapeutic tool to combat microbial resistance. Advanced Drug Delivery Reviews. 2013;(65):1803– 1815. 10.1016/j.addr.2013.07.011.

